# Cocktail Chemical Labeling for In-Depth Surfaceome Profiling of Bone-Marrow Derived Dendritic Cells

**DOI:** 10.1101/2025.03.15.643439

**Authors:** Shiming Sun, Siyang Liu, Tao Deng, Yudi Hu, Rong Yang, Jie P. Li, Yun Ge

**Affiliations:** Institute of Chemical Biology, Shenzhen Bay Laboratory, Shenzhen 518132, China; State Key Laboratory of Coordination Chemistry, Chemistry and Biomedicine Innovation Center (ChemBIC), School of Chemistry and Chemical Engineering, Nanjing University, Nanjing 210023, China; Department of Urology, Nanjing Drum Tower Hospital, Affiliated Hospital of Medical School, Nanjing University, Nanjing 210008, China; State Key Laboratory of Chemical Oncogenomics, School of Chemical Biology and Biotechnology, Peking University Shenzhen Graduate School, Shenzhen 518055, China

**Keywords:** cell surface proteomics, ruthenium complex, photoredox catalysis, dendritic cells

## Abstract

Cell surface proteins (CSPs) are crucial identifiers for cell types and states, especially in dendritic cells (DCs). Current proteomic methods for profiling CSPs are limited by their hydrophobic nature and low abundance, which often require genetic or cell surface engineering and exhibit biased and insufficient labeling efficiency. Herein, we report [Ru(bpy)_3_]Cl_2_ (Ru) for effective biotinylation on the cell surfaceome via a simple “mix and lighten” method. The versatile photoredox pathways of Ru are leveraged using a probe cocktail of biotin-phenol and biotin-hydrazide for improved substrate coverage. The “cocktail” labeling strategy results in reproducible identification of up to 733 plasma membrane proteins on HeLa cells, and is further applied to map dynamic changes in the surfaceome during the differentiation of primary bone marrow-derived dendritic cells, which demonstrates a user-friendly and deep-surfaceome-coverage tool for profiling dynamic changes in primary cells, with potential implications for cell identities, functional states, and novel drug targets.

## 1. Introduction

The variable proteins on the cell surface, collectively termed the surfaceome, functionally engage in many pivotal biological processes, spanning from interactions with the neighbors and microenvironments to signal transduction.^[1]^ This makes them valuable markers for distinguishing cell types and states, as well as promising targets for drug discovery.^[2]^ For instance, cells in the immune system are typically immunophenotyped according to their cell surface proteins (CSPs) using the cluster of differentiation (CD). Dendritic cells (DCs), key players in the immune system, are identified by a panel of diverse co-existing CD molecules on their surface.^[3]^ These cells capture environmental signals and present them to other immune cells, coordinating both innate and adaptive immune responses. The diverse cell surface proteotypes enable classification of different DC subtypes, linking them to their identity and function. However, these validated CD molecules represent only a small fraction of CSPs, hindering the discovery of new cell identifiers and functional players responsive to complex environments. Although single-cell sequencing offers insights into cell types at the transcriptomic level, these results do not always align well with bona fide protein expressions. Therefore, comprehensive surfaceome profiling remains to be the long-lasting demand for understanding cell identities, functional states, and for discovering novel drug targets.

Mapping CSPs is primarily done through mass spectrometry-based proteomic analyses. However, due to the hydrophobic nature and low abundance of CSPs, enrichment or separation steps are often required for high-coverage identification.^[1a]^ The strategy of direct cell surface protein biotinylation, followed by affinity purification, is widely used for surfaceome profiling. We reason that an ideal biotinylation strategy should be: 1) simple and rapid, 2) compatible with proteomic analyses, 3) label without requiring additional engineering, 4) with high labeling efficient and protein coverage, and 5) adaptable for primary cells and tissues. Achieving these criteria would facilitate comprehensive surfaceome profiling while maintaining cellular integrity and physiological relevance. Representative biotinylation methods include sulfo-NHS-SS-biotin and the cell surface capture (CSC) technique.^[4]^ However, sulfo-NHS-SS-biotin labeling is biased towards lysine residues, leading to possibly reduced trypsin cleavage efficiency; CSC requires multiple steps and suffers from reproducibility issues due to natural glycosylation heterogeneity across cell types and physiological states. These challenges have driven the development of more efficient chemical tools, such as peroxidase-based catalysts (e.g., APEX2, HRP), which have been widely used to map dynamic surfaceome changes in various biological contexts, including oncogene transformation and neuronal development.^[5]^ However, these tools generally require membrane-tethering for enhanced labeling efficiency^[5c]^ and generate phenol radicals predominantly reacting with tyrosine, resulting in biased labeling towards Tyr-rich membrane proteins.^[6]^ Therefore, more versatile and efficient labeling methods are needed to improve usability and overall CSP coverage.

In this study, we present a simple, fast, and efficient cocktail chemical labeling strategy for in-depth surfaceome profiling. Unlike conventional cell surface-tethered biotinylators, we discovered that free Ru is highly effective for labeling CSPs using biotin hydrazide (BHy) or biotin phenol (BP), achieving optimal labeling efficiency within just 2 minutes. When integrated with proteomic analysis, both probes resulted in greater identification of plasma membrane proteins (PMPs) compared to a previously reported soluble HRP-mediated biotinylation strategy, PECSL.^[5f]^ By exploiting two Ru-catalytic pathways, we developed a probe cocktail containing both BP and BHy, which significantly increased the number of identified surfaceome proteins with high sequence coverage. Using this cocktail chemical labeling strategy, we systematically profiled CSP dynamics during the differentiation and maturation of primary bone marrow-derived dendritic cells (BMDCs), linking surfaceome changes to different cellular stages. This work provides, for the first time, a comprehensive map of sufaceome dynamic changes during BMDC differentiation, offering new insights into putative new surface markers and the underlying molecular mechanisms. The simple “mix and lighten” method, utilizing commercially available Ru complexes and biotin probe cocktails, shows great promise for mapping dynamic surfaceomes in primary cells and even complex patient-derived clinical samples, with comprehensive coverage.^[5f]^

### 2. Results

### 2.1. Establishment of Ru-mediated photocatalytic reaction for CSP labeling

Transition metal complexe (TMC)-based photosensitizers exhibit high photocatalytic efficiency due to their ability to engage in multiple catalytic pathways.^[7]^ We hypothesized that a single catalyst could efficiently label cell surface proteins (CSPs), improving surfaceome coverage and reducing labeling bias through multiple mechanisms. Recently, we identified the blue-light-activated photocatalyst [Ru(bpy)_3_]Cl_2_ (Ru) as a bifunctional and highly efficient catalyst for capturing cell-cell interactions. This catalyst efficiently converts the biotin-phenol probe into phenoxy radicals via the single-electron transfer (SET) process.^[8]^ Encouraged by these properties, we sought to repurpose Ru for surfaceome labeling (**Figure S1A**) to enhance labeling efficiency and surfaceome coverage.^[8]^ Previous studies have shown that cell-tethered HRP and APEX2 are more effective for surfaceome labeling than free enzymes. To determine if similar effects would occur with Ru, we explored three non-invasive methods for tethering Ru to the cell membrane, targeting different membrane layers: (1) the fucosyl derivative of Ru (GDP-Fucose-Ru, abbreviated as Ru-G) that can be transferred to LacNAc units on N-glycoproteins via 1,3-fucosyltransferase; (2) NHS ester-functionalized Ru (abbreviated as Ru-N) that can react with active amines on lysine residues of cell membrane proteins; and (3) the 1,2-dipalmitoyl-sn-glycero-3-phosphoethanolamine (DPPE)-conjugated Ru (abbreviated as Ru-L) that can insert into the lipid bilayer via hydrophobic interactions (**Figure 1A**). Surprisingly, while the surface labeling efficiency of these Ru conjugates partially correlated with their loading amounts (**Figure S1B**), none achieved a similar biotinylation level to soluble free Ru (**Figure 1B**). In contrast, cell surface tethering of other catalysts, including HRP and the photosensitizers DBF and Ir (**Figure S1C, S1D and S1E**), resulted in higher biotinylation intensities compared to their soluble forms (**Figure 1C**). Notably, free Ru exhibited the highest labeling efficiency with the shortest labeling time, highlighting its unique and robust capacity for promiscuous surface biotinylation. We also confirmed that free Ru demonstrated minimal spontaneous penetration or cellular uptake, as shown by intracellular Ru signals measured after 60 minutes of incubation (**Figure 1D**), suggesting that the Ru-catalyzed reactions occurred exclusively on the cell surface. These findings prompted us to optimize the surface biotinylation process using free Ru as the catalyst.

**Figure 1.**
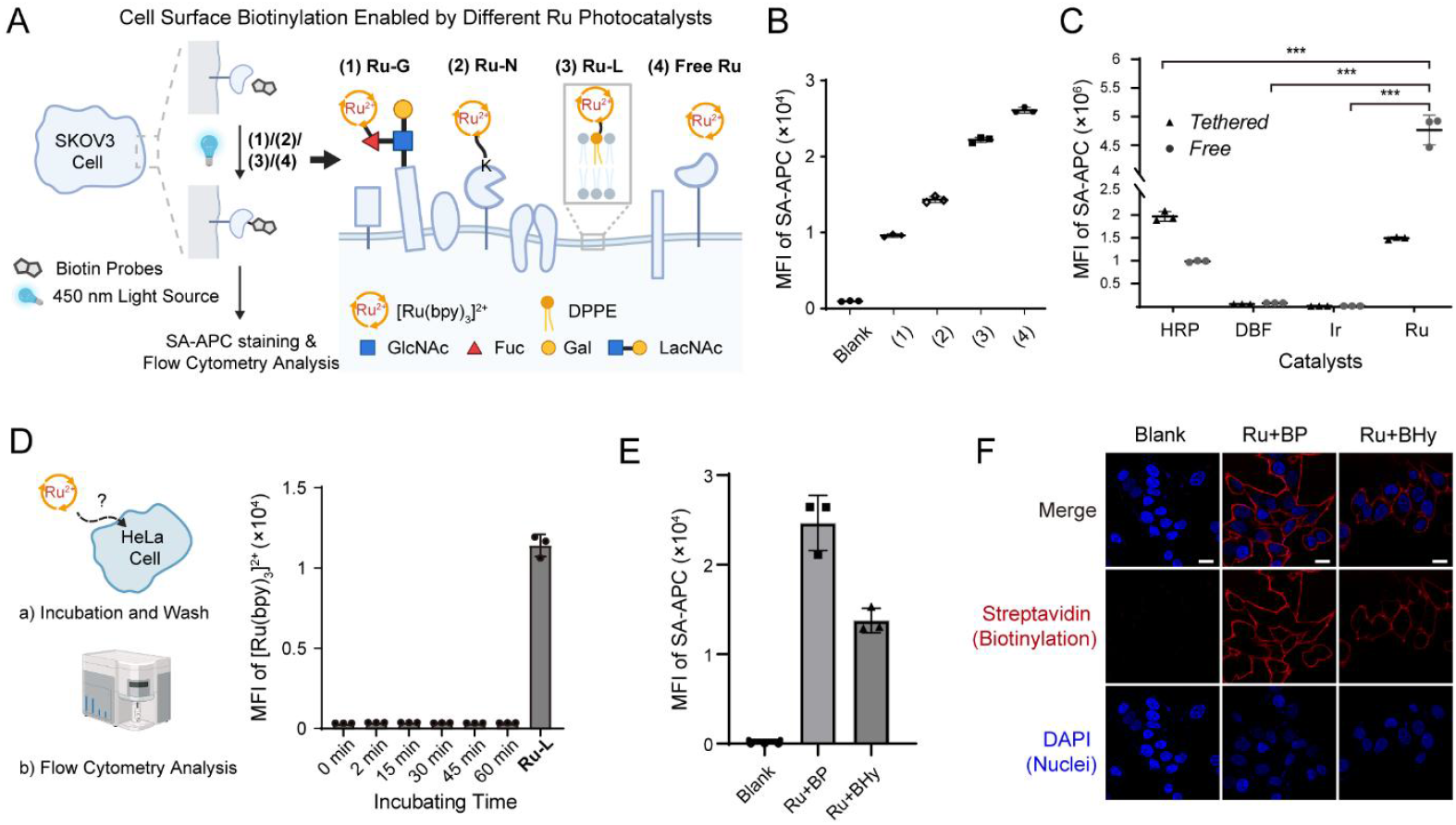
Free [Ru(bpy)_3_]Cl_2_ (Ru) photocatalyst enables highly efficient cell surface protein biotinylation. (A) Schematic depiction of cell surface biotinylation screening of different Ru formats using SKOV3 cells. (1) Ru-G, GDP-Fucose-Ru; (2) Ru-N, NHS ester-functionalized Ru; (3) Ru-L, DPPE-conjugated Ru; (4) free Ru catalyst. (1)-(3) were tethered on the cell membrane and (4) were free in the medium. All labeling reactions with the addition of the biotin phenol probe (BP) and APS were initiated by irradiation with a 450 nm light source. (B) Comparison of cell surface biotinylation among (1)-(4). (C) Comparison of cell surface biotinylation using free and tethered formats of HRP, DBF, Ir and Ru. The one-way ANOVA test was used for statistical analysis. (D) Flow cytometry analysis of cellular uptake of the free Ru catalyst. HeLa cells were incubated with free Ru for the indicated time; Ru-L was selected as the positive control. (E) Cell surface biotinylation of Ru-mediated labeling using BP and biotin hydrazide (BHy). Data in (B), (C), (D) and (E) are represented as mean ± S.D. (n=3) (F) Confocal microscopic images of HeLa cells after free Ru-catalyzed cell surface biotinylation. Representative images of two biological replicates were shown. Scale bar: 10 μm.

Ru catalysts have been reported to activate BP or BHy through two distinct mechanisms: energy transfer and SET pathways. First, we found that adding APS enhanced the labeling intensity of both Ru+BP and Ru+Bhy (**Figure S2A**). Cell viability remained above 70% even after 60 minutes when Ru concentrations were below 10 μM and in the absence of probes (**Figure S2B**). Based on these observations, we optimized the Ru concentration to 10 μM in the presence of 200 μM BP probe, 2 mM APS, and 5 minutes of light exposure (**Figure S2C**). Assessment of light exposure time revealed a 2-minute light exposure sufficient for saturated labeling (**Figure S2D**). Next, we proceeded to refine probe concentrations under these optimal conditions. A 200 μM BP concentration yielded maximal cell labeling (**Figure S2E and S2F**), whereas BHy exhibited a positive correlation between probe concentration and biotinylation intensity, with 500 μM BHy providing substantial surface labeling (**Figure S2G and S2H**). Both probes showed negligible cytotoxicity in multiple immune cell lines (**Figure S2I**), and achieved remarkable labeling intensities, with the Ru+BP method outperforming the Ru+BHy method in labeling efficiency (**Figure 1E**). Immunofluorescence imaging of biotinylated proteins further confirmed that free Ru-catalyzed biotinylation exclusively occurred on cell surfaces (**Figure 1F**). These results demonstrate that free Ru with 2-minute blue light irradiation enables specific, high-efficiency cell surface protein biotinylation, making it a promising tool for profiling the cell surface proteome with excellent biocompatibility.

### 2.2. Substrate scope of the Ru+BHy labeling is different from that of the Ru+BP labeling

Building on the optimized free Ru-catalyzed surface labeling strategy, we next integrated the labeling reaction with the affinity purification mass spectrometry (AP-MS) workflow. Given that the PECSL method using exogenous HRP and the BP probe in the presence of H_2_O_2_ has been reported for rapid and efficient surfaceome profiling,^[5f]^ we compared our Ru-catalyzed surfaceome profiling strategy with PECSL (hereafter referred to as HRP). Briefly, living cells were resuspended in PBS buffer and incubated with Ru, APS and biotin probes. The labeling reaction was initiated by blue light irradiation for 2 minutes. Afterward, cells were lysed, and the biotinylated proteins were enriched using streptavidin resin, trypsinized into peptides, and analyzed by mass spectrometry (**Figure 2A**). For comparison, HRP-based surfaceome labeling was performed in parallel, with two replicates conducted for each condition to ensure high sequence coverage reproducibility and generate overlapping groups for subsequent analysis (**Figure S3A and S3B**). The identified proteins were annotated using the UniProt Gene Ontology Cellular Component (GOCC) database, with particular focus on plasma membrane proteins (PMPs) and cell surface proteins (CSPs) based on a previous method.^[4c]^

**Figure 2.**
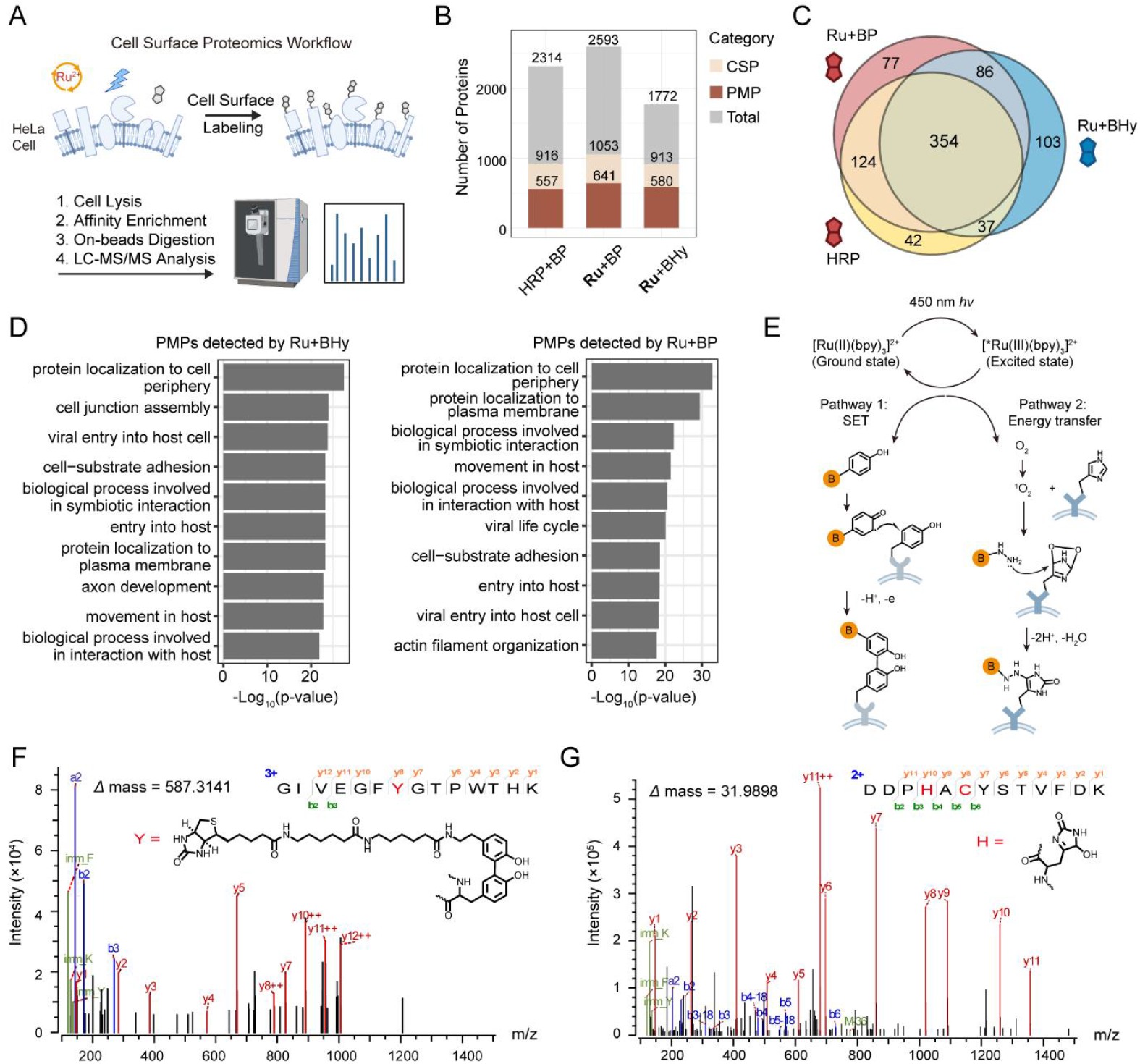
Comparison of surfaceome profiling using two Ru-mediated labeling approaches, Ru+BHy and Ru+BP. (A) Workflow of Ru-mediated cell surface labeling and mass spectrometric analysis. (B) Number of plasma membrane proteins (PMPs) and cell surface proteins (CSPs) detected by different cell surface labeling methods. CSPs and PMPs were annotated according to GOCC database. Results are the intersection of two biological replicates. (C) Venn diagram demonstrating identified PMPs from three labeling methods. (D) Comparison of enriched Gene Ontology Biological Process (GOBP) terms of PMPs detected only by Ru+BHy and Ru+BP, respectively. (E) The two distinct labeling pathways of Ru-catalyzed cell surface biotinylation using BP and BHy probes. Representative MS/MS spectra of (F) BP probe-modified tyrosine residue of *Cp*OGA^D298N^ and (G) dioxidized histidine residue of BSA peptide.

Using this workflow, we identified a total of 916 CSPs and 557 PMPs for HRP, 913 CSPs and 580 PMPs for Ru+BHy, and 1053 CSPs and 641 PMPs for Ru+BP, respectively. Both Ru-catalyzed methods slightly outperformed HRP (**Figure 2B**). We focused on PMPs, which were annotated using a more stringent PMP-based strategy to provide more accurate and physiologically relevant insights into cell states. While all three methods detected the major 354 PMPs, both Ru+BHy and Ru+BP methods identified additional unique PMPs not detected by HRP (**Figure 2C**). These unique proteins also possessed higher sequence coverage, suggesting the superior labeling efficiency of the Ru-based approaches (**Figure S3C**). Although there was significant overlap in the PMPs identified by both Ru probes, there were also differences, resulting in enriched Gene Ontology Biological Process (GOBP) terms specific to each labeling strategy (**Figure 2D**).

We have previously shown that the blue light-activated Ru catalyst can engage in two energy transfer pathways (**Figure 2E**): the SET pathway generating highly reactive phenoxyl radicals for BP-tyrosine conjugation; another energy transfer pathway producing oxidized histidine (His) for BHy attachment.^[8]^ Accordingly, we hypothesized that differences in labeling efficiency and substrate bias may arise from the distinct mechanistic preferences of the biotin probes for different amino acid residues. To test this, we performed Ru-catalyzed labeling on model proteins (bovine serum albumin, BSA, and *Clostridium perfringens* O-GlcNAcase mutant, *Cp*OGA^D298N^) using BP or BHy, and detected possible chemical modifications by mass spectrometry (**Figure S3D**). The BP-Tyr conjugate showed a mass shift of 587.3141 Da on Tyr in the *Cp*OGA^D298N^ sample treated with Ru+BP. While the BHy adduct was not observed, likely due to its unstable chemical properties and suboptimal ionization efficiency, a notable mass shift of 31.9908 Da corresponding to oxidized His was observed in the ‘Ru+BHy’-treated BSA (**Figure 2F**). These results further confirm the two distinct labeling mechanisms, which may explain the high efficiency and the discrepancy observed in Ru-catalyzed BP/BHy labeling on cell surfaces.

### 2.3. Probe cocktail labeling afforded deeper coverage of the surfaceome

Interestingly, mass spectrometry analysis of ‘Ru+BP’ labeling on *Cp*OGA^D298N^ also revealed the dioxidized histidine modification, indicating that ^1^O_2_ may be produced in the Ru+BP approach as well (**Figure S3E**). The coexistence of these two mechanisms encouraged us to combine two probes into a cocktail chemical labeling strategy for deeper surfaceome profiling (**Figure 3A**). Based on the optimization results from single-probe Ru-catalyzed labeling, we prepared a probe cocktail containing 200 µM BP and 500 µM BHy for cell surface labeling, with 10 µM Ru and 2 mM APS. The surface labeling was performed under 2 minutes of blue light irradiation, showing minimal cytotoxicity comparable to other labeling methods (**Figure S4**). Both flow cytometry and immunoblotting analyses indicated that the cocktail achieved the highest labeling efficiency—1.2 to 1.5 times that of Ru+BP and 2 to 5 times that of Ru+BHy (**Figure 3B and 3C**). Consistent with these results, LC-MS/MS analyses further confirmed the superior efficacy of the cocktail, leading to a significant increase in both identified CSPs and PMPs (**Figure 3D; Supplementary Table 1**). Of note, the cocktail chemical labeling method identified 92 and 153 additional PMPs compared to Ru+BP and Ru+BHy, respectively. Moreover, the PMPs detected by the cocktail closely resembled the combined set of PMPs identified by BP- and BHy-mediated labeling (**Figure 3D**). Since the use of two probes targets more available residues for subsequent enrichment and MS detection, we analyzed proteins detected by all three Ru-based methods and confirmed that the cocktail approach provided the highest sequence coverage overall (**Figure 3E**). These results suggest that Ru-catalyzed cocktail labeling increases surfaceome coverage by integrating two distinct substrate targeting mechanisms.

**Figure 3.**
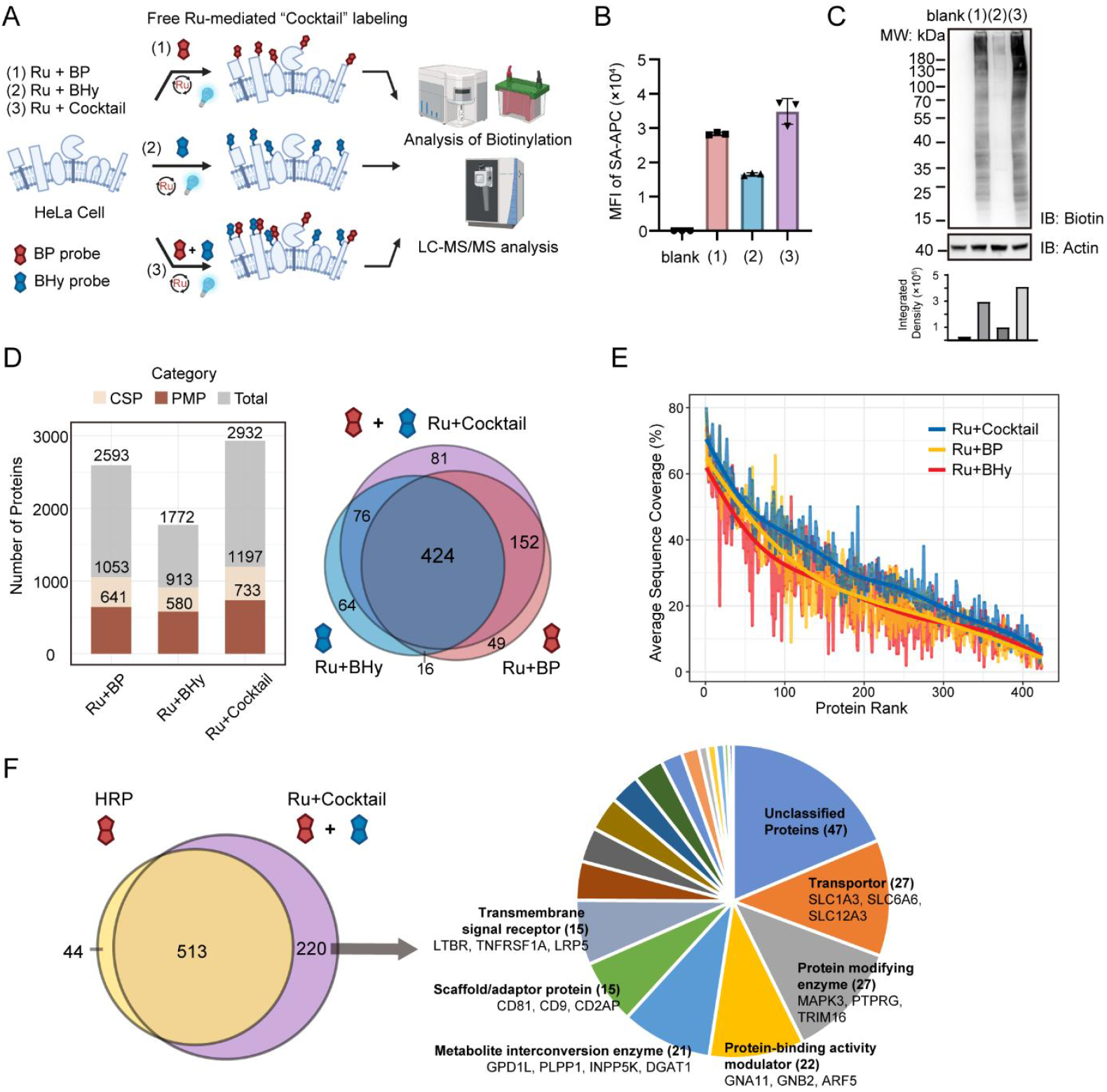
Ru-catalyzed cocktail chemical labeling yielded higher cell surface protein labeling efficacy and higher coverage. (A) Scheme of Ru-catalyzed labeling workflows using a single probe or the probe cocktail for comparision. Flow cytometry (B) and immunoblotting analyses (C) demonstrated that a probe cocktail resulted in the highest labeling intensity. (D) Histogram and Venn diagram showing comparisons of identified proteome using Ru+Cocktail with those using Ru+BHy and Ru+BP. (E) Comparison of average sequence coverage of PMPs reproducibly identified in two replicates by the three methods. (F) The number of PMPs identified by Ru-catalyzed cocktail chemical labeling compared to HRP was shown. Representative proteins and their types exclusively detected by the cocktail labeling were annotated.

Finally, we compared the PMPs identified by the cocktail labeling method with those obtained using the HRP method. The cocktail labeling technique identified 220 additional unique PMPs, representing a nearly 40% increase (**Figure 3F**). This group included PMPs with crucial cellular functions: transporters such as SLC1A3, SLC6A6, SLC12A3; transmembrane signal receptor like LTBR, TNFRSF1A and LRP5; scaffold/adaptor proteins such as CD81, CD9 and CD2AP; proteins involved in protein-binding activities like GNA11, GNB2, and ARF5; and membrane-residing enzymes like TRIM16 and PTPRG. This additional set of PMPs enabled by the cocktail strategy were envisioned to advance our understandings of related key biological processes. Together, we demonstrated that the Ru-catalyzed cocktail chemical labeling approach greatly expands the surfaceome landscape due to its fast kinetics, high efficiency, and broad coverage, which was expected to provide more new surface marker candidates and insights into cell states.

### 2.4. The Ru+Cocktail approach enabled primary BMDCs surfaceome profiling

The robustness, convenience, low cytotoxicity, and, most importantly, the lack of any need for cell engineering make the Ru+Cocktail approach well-suited for surfaceome profiling in primary cell samples. Therefore, we focused on mouse primary BMDCs. Dendritic cells (DCs) are considered one of the most important antigen-presenting cells (APCs) due to their ability to initiate immune responses by activating naïve T cells and coordinating innate and adaptive immunity.^[3]^ During this highly regulated process, DCs undergo profound phenotypic and functional reprogramming, which alters their interactions with the environment and other cells.^[9]^ Traditionally, this process has been described using a limited set of DC-specific markers, offering only a fragmented view of the broader dynamics.^[10]^ Recent advances in DC biology have highlighted the extensive role of PMPs,^[11]^ spurring efforts to better understand the surfaceome during key biological processes. However, to our knowledge, previous proteomic studies of mouse BMDCs have provided limited information, with only 192 surface proteins identified.^[12]^ We believe that our cocktail labeling method could explore surfaceome dynamics during the differentiation of mouse primary BMDCs (**Figure 4A**), offering a comprehensive atlas of these changes.

**Figure 4.**
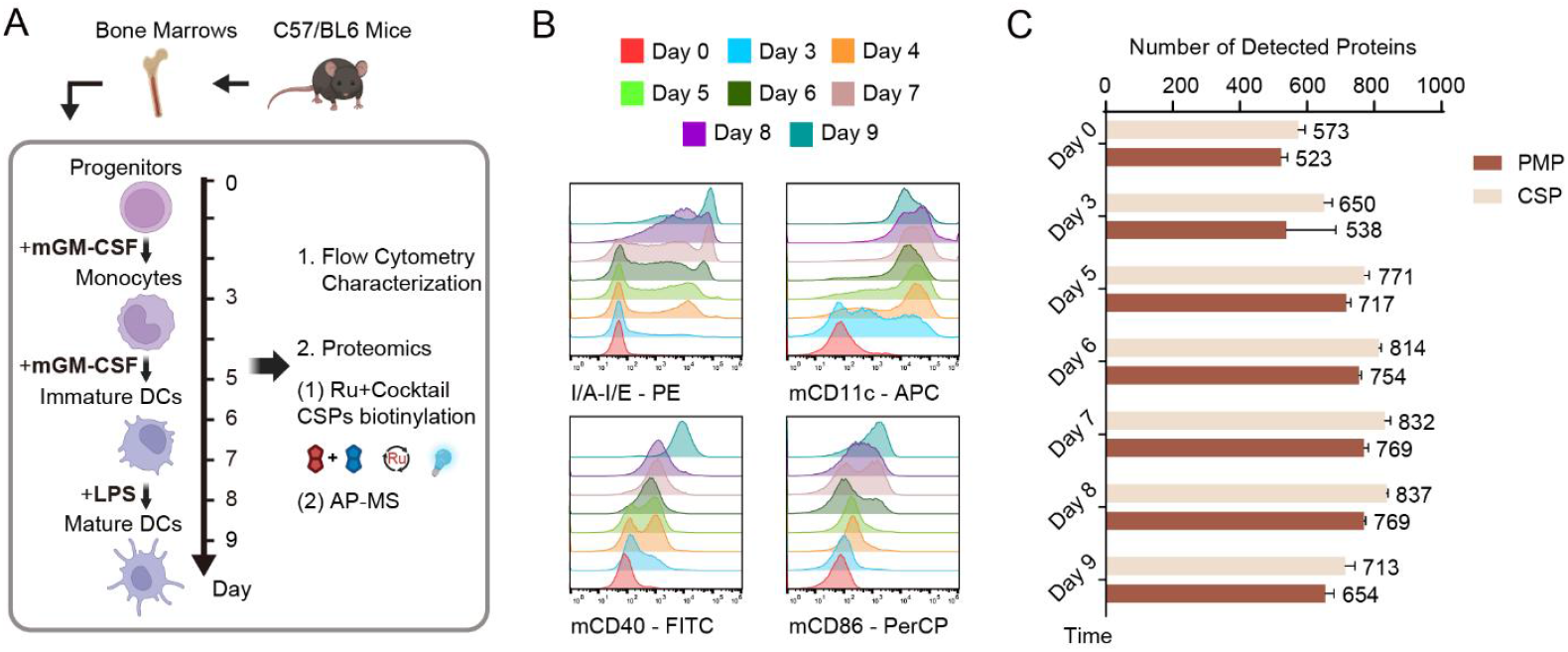
Cell surface proteome profiling during differentiation and maturation of BMDCs. (A) Schematic workflow of surfaceome profiling during BMDCs’ differentiation and maturation. Bone marrows of C57/BL6 mice were harvested for BMDC generation in vitro under stimulation of mGM-CSF. At the end of Day 8, immature BMDCs were treated with LPS to produce mature BMDCs. During these processes, cell samples were collected and submitted to flow cytometry analyses or cocktail chemical labeling. (B) Flow cytometry analyses of BMDC samples at indicated time points in (A) using indicated known surface markers. (C) Numbers of cell surface annotated proteins (CSPs), and plasma membrane annotated proteins (PMPs) detected in indicated samples. Data are represented as mean ± S.D (n=4).

We employed an in vitro differentiation model that includes successive cellular stages under the stimulation of mouse granulocyte-macrophage colony-stimulating factor (mGM-CSF).^[13]^ Previous studies of BMDC generation and maturation have shown gradual expression of surface antigen-presentation molecules and co-stimulatory receptors (e.g., MHC class II, CD80, CD86), enhancing the DCs’ ability to interact with T cells and activate T cell-mediated immune responses.^[14]^ To validate the differentiation process, we first performed flow cytometry analysis of known surface markers, including I-A/I-E molecules and CD11c (**Figure 4B and S5A**).

Next, primary BMDCs from days 3, 5, 6, 7, 8, and 9 were collected and subjected to cocktail chemical labeling for surfaceome profiling. We were pleased to find that Ru-catalyzed CSP biotinylation was as efficient as in cell line studies (**Figure S5B**). In total, we identified 600 to 800 PMPs for each condition, with nearly 1200 PMPs detected across all samples (**Figure 4C; Supplementary Table 2)**. The number of PMPs/CSPs detected during differentiation showed an initial increase followed by a subsequent decrease. Pearson correlation and principal component analysis (PCA) revealed high consistency among samples collected on the same day (**Figure S5C and S5D**). More importantly, the PCA plot effectively illustrated the continuous slow changes and rapid changes in PMP patterns during the processes of differentiation and maturation, consistent with our understanding of BMDC differentiation. As a result, we divided the 10-day process into two phases for analysis. From Day 0 to Day 8, progenitor cells differentiate into immature DCs under mGM-CSF stimulation, which was ideal for analyzing surfaceome changes during BMDC differentiation. The second phase, between Day 8 and Day 9, was marked by a profound proteomic reshaping triggered by lipopolysaccharide (LPS) stimulation, leading to mature DCs. In conclusion, these promising results underscored the applicability of Ru-mediated surfaceome profiling for primary immune cells. Unlike previous studies that relied on a limited set of surface markers, the dataset generated here provides the most comprehensive surfaceome dynamics of BMDC generation to date.

### 2.5. Investigation of cell surface proteome dynamics during BMDCs’ maturation

At last, we analyzed this dataset to identify previously undocumented changes in surface proteins during BMDC differentiation and maturation using label-free quantification (LFQ). To validate the reliability of our surfaceomic data, we first compared the correlations between flow cytometric and proteomic results. Quantitative data of several known key maturity markers (MHC II, mCD11c, mCD86, mCD40) all showed high correlation with previous flow cytometric data (**Figure S6A and S6B**). Next, in order to investigate PMP expression profiles of BMDCs, we categorized all surface proteins identified during the first BMDC differentiation phase (Days 3 to 8) using an unsupervised fuzzy clustering approach (**Supplementary Table 3**), which resulted in six distinct clusters based on surface abundance profiles (**Figure 5A**).^[15]^ Cluster 2 proteins exhibited a steep early increase, while clusters 1, 4, and 6 showed dynamic changes. Notably, clusters 3 and 5 contained proteins that decreased or increased stepwise over these days, respectively (**Figure 5A**). We focused on proteins in clusters 3 and 5 to determine if these proteins could reflect functional dynamics during specific stages of differentiation (**Figure 5B and 5C**). GOBP terms specifically enriched in cluster 5 were associated with endocytosis/phagocytosis and cytokine production, indicating increased immune activity of BMDCs during differentiation and maturation.^[16]^ For cluster 3, we found enrichment in terms related to cell adhesion, consistent with the observation that BMDCs increase mobility during maturation to migrate to distal regions.^[17]^ These results support the theory that BMDC progenitors in the bone marrow give rise to circulating precursors that home to tissues, where they reside as immature DCs with high phagocytic capacity.^[18]^ Unlike previous antibody-based methods (e.g., flow/mass cytometry), the Ru-labeling approach revealed these dynamic surface-abundance profiles without prior knowledge, highlighting stage-specific functional dynamics in BMDC maturation. We also validated these surfaceome dynamics during the first phase of BMDC differentiation using standard flow cytometry. We identified numerous CD molecules in clusters 3 and 5 (**Figure S7; Supplementary Table 4**). Among these, we focused on CCR2 and CCR7 representing chemokine receptor expression levels, CD36 indicating responses to DAMPs and PAMPs,^[19]^ as well as the previously underrepresented CD270 (HVEM). Flow cytometry data showed a strong correlation with mass spectrometric results for these markers (**Figure 5D**), further confirming the high fidelity of this Ru+Cocktail-based surfaceome mapping approach. Among these markers, HVEM has been reported to play a role in immune regulation and inflammatory responses through ligand binding,^[20]^ suggesting that upregulating HVEM could influence immune response modulation, DC differentiation, and other functions.

**Figure 5.**
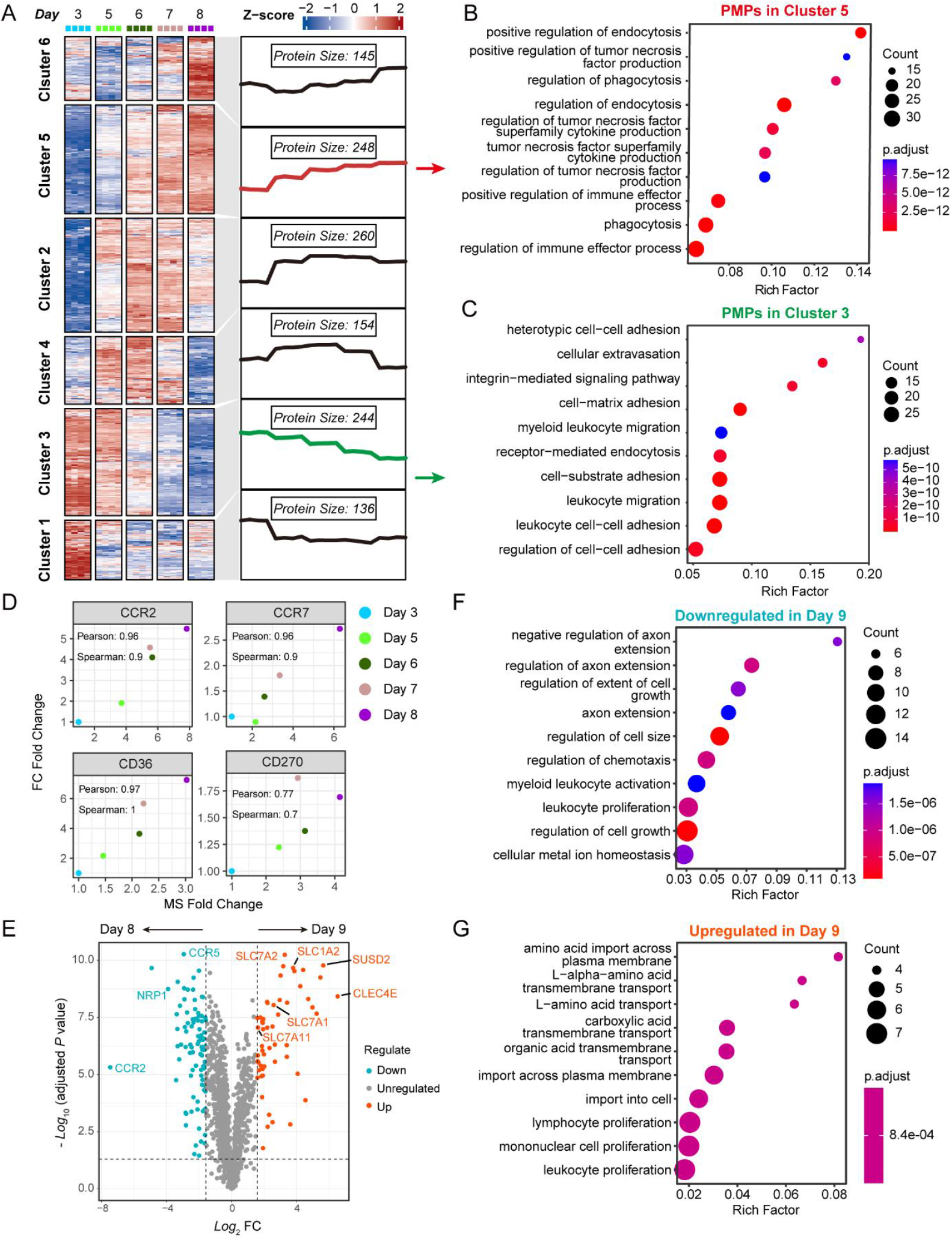
Surfaceome abundance profiles reveal dynamic patterns during BMDC differentiation. (A) Unsupervised c-means fuzzy clustering of surfaceome abundance profiles reveals six distinct clusters. Gene ontology terms overrepresented in biological processes for (B) the stepwise upregulated cluster, cluster 5, and (C) the stepwise downregulated cluster, cluster 3. (D) Correlation analysis of expression dynamics between mass spectrometry (MS)- and flow cytometry (FC)-determined fold change for the selected CD antigens using mean value of all experiments. “Pearson” and “Spearman” stand for Pearson’s correlation and Spearman’s correlation, respectively. (E) Volcano plot shows the surfaceome quantification comparison of immature BMDCs versus mature BMDCs. Adjusted P value = 0.05 and ± 3-fold change (FC) are denoted by gray dashed lines as the significance threshold. GOBP terms enriched in the downregulated (F) and upregulated (G) PMPs on Day 9 versus Day 8.

We then investigated the second phase of BMDC maturation with LPS stimulation, which was expected to involve dramatic changes in the surfaceome and cellular functions. Differential expression analysis between the surfaceome of Day 9 and Day 8 revealed downregulated proteins associated with cell adhesion and morphology regulation, including CCR2, CCR5, and NRP1, suggesting a transition towards a more stable morphological state following LPS stimulation (**Figure 5E, 5F and Supplementary Table 5**). Upregulated proteins were linked to amino acid transport (**Figure 5G**), including SLC1A2, SLC7A1, SLC7A2, and SLC7A11, suggesting increased amino acid uptake by BMDCs (**Figure 5E**). Previous studies have indicated that SLC proteins involved in amino acid metabolism may influence IFN-I production, redox balance, antigen presentation, and cellular metabolism in DCs.^[11b]^ The upregulation of these amino acid transport-related SLC proteins may represent potential targets on mature DCs.

We also looked into these upregulated proteins between Day 9 and Day 8. CLEC4E (mincle) is classified as a group II C-type lectin-like receptor expressed on myeloid DCs, monocytes and macrophages.^[21]^ Our data align with previous transcriptome-level findings.^[22]^ CLEC4E has been shown to induce the production of TNF, IL-10, and CXCL2. Moreover, a homologous protein, CLEC9A, has been used as a DC marker in designing a DC-T cell engager.^[23]^ These findings suggest that CLEC4E could be an important target for modulating DC function or phenotyping.

## 3. Conclusion

In summary, we have developed a Ru-catalyzed “cocktail” labeling strategy for surfaceome profiling with high coverage when integrating with proteomic analysis. Intriguingly, we observed that the high solubility and membrane-impermeant properties of [Ru(bpy)_3_]Cl_2_ resulted in higher labeling efficiency compared to its membrane-tethered form. By utilizing readily available free-state Ru, combined with biotin probes and a 2-minute blue light irradiation, we eliminate the need for genetic or surface modification, as well as the cytotoxic H_2_O_2_, simplifying the entire labeling process. The dual mechanisms of Ru-catalyzed photoredox reactions inspired us to introduce a ‘cocktail’ labeling strategy to achieve higher protein identification coverage. We demonstrated that this approach significantly increased the number of identified CSPs by expanding the labeling residues, resulting in approximately a 40% increase compared to the HRP-mediated surface biotinylation method, PECSL. This ‘cocktail’ strategy is also potentially applicable to various metal photocatalysts and enzymes, enabling multiple labeling mechanisms in a single reaction for expanded substrate coverage. In addition to its high efficiency and expanded labeling sites, TMC-catalyzed photo-reaction allows customization of the labeling radius for tunable spatial precision. This work presents a novel application of Ru-catalyzed protein labeling for surfaceome profiling, complementing previous applications for detecting cell-cell interactions and profiling protein-specific microenvironments. Future efforts can focus on optimizing sample preparation processes and applying a ratiometric approach to improve spatial specificity in proteomic results.^[24]^

From a biological perspective, our investigation of the differentiation and maturation processes of BMDCs has resulted in a comprehensive atlas of dynamic surfaceome changes, the first of its kind to our knowledge. This atlas offers valuable biological insights into DCs from a surfaceome proteotype perspective, potentially offering new ways to define how DCs detect pathogens or pathological changes and distinguishing different functional states of DCs. Specifically, we speculate that proteins exhibiting substantial variations in abundance during differentiation and maturation may play key roles and warrant further investigation into their underlying molecular mechanisms. Additionally, the success of profiling the dynamic surfaceome of DCs has motivated us to extend this strategy to other cell types or clinical samples.

In conclusion, we reported a Ru-catalyzed cocktail labeling strategy and demonstrated its applicability on profiling dynamic surfaceome changes during DC maturation. We anticipate that this method will find widespread applications in accurately classifying cell subtypes with distinct functionalities, tracking cellular evolution at the proteotype level, and identifying potential biomarkers and drug targets, thereby complementing transcriptomic profiling results.

## Supporting Information

Supplementary Figures.

Supplementary Table 1. Protein identification results of different cell surface labeling approaches including Ru+BP, Ru+BHy, Ru+Cocktail and HRP-based technique (xlsx) Supplementary Table 2. Protein identification results for BMDC samples from Day 0 to Day 9 using the Ru+Cocktail labeling approach (xlsx)

Supplementary Table 3. LFQ results for BMDC samples from Day 0 to Day 9 using the Ru+Cocktail labeling ap-proach and time-series clustering annotations for Day 3 to Day 8 results (xlsx)

Supplementary Table 4. CD molecules identified in BMDC surfaceome LFQ results (xlsx) Supplementary Table 5. Differential expression analysis be-tween Day 9 and Day 8 using log2 transformed LFQ results from Supplementary Table 3 (xlsx)

## Acknowledgements

This research was supported by the National Key R&D Pro-gram of China (2019YFA09006600 to J.P.L., 2024YFC3407200 to Y.G.), the National Natural Science Foundation of China (22277080 and 92478103 to Y.G., 21977048 and 92053111 to J.P.L.), the Fundamental Research Funds for the Central Universities (J.P.L.), Natural Science Foundation of Jiangsu Province (BK20202004 to J.P.L.), Beijing National Laboratory for Molecular Sciences (BNLMS202008 to J.P.L.), Program for Innovative Talents and Entrepreneur in Jiangsu (J.P.L.), and the Excellent Research Program of Nanjing University (ZYJH004 to J.P.L.), Major Programs of Shenzhen Bay Laboratory and Shenzhen Bay Laboratory Startup Fund (S241101001, S211101001 and 21230102 to Y.G.). Technical support from the Shenzhen Bay Laboratory Core Facility, especially bioimaging, proteomics and animal facilities, is gratefully acknowledged.

## Author Contributions

Y.G. and J.P.L. designed the experimental strategies. S.S. performed the experiments with the help of S.L., Y.H., and T.D.. S.S., Y.G. and J.P.L. prepared the figures. The manuscript was written by S.S., Y.G. and J.P.L. with approval of all authors.

## Data Availability Statement

The mass spectrometry data generated in this study have been deposited to the ProteomeXchange Consortium via the iProX partner repository with the dataset identifier PXD060940.^[26]^

## References

[1] a) C. C. Wu, J. R. Yates III, Nat. Biotechnol. 2003, 21, 262–267; b) M. S. Almén, K. J. Nordström, R. Fredriksson, H. B. Schiöth, BMC Biol. 2009, 7, 50; c) D. Bausch-Fluck, U. Goldmann, S. Müller, M. van Oostrum, M. Müller, O. T. Schubert, B. Wollscheid, Proc. Natl. Acad. Sci. U. S. A. 2018, 115, E10988–E10997.

[2] a) G. P. Adams, L. M. Weiner, Nat. Biotechnol. 2005, 23, 1147–1157; b) F. E. Craig, K. A. Foon, Blood 2008, 111, 3941–3967; c) Z. Hu, J. Yuan, M. Long, J. Jiang, Y. Zhang, T. Zhang, M. Xu, Y. Fan, J. L. Tanyi, K. T. Montone, O. Tavana, H. M. Chan, X. Hu, R. H. Vonderheide, L. Zhang, Nat. Cancer 2021, 2, 1406–1422.

[3] a) J. Banchereau, F. Briere, C. Caux, J. Davoust, S. Lebecque, Y.-J. Liu, B. Pulendran, K. Palucka, Annu. Rev. Immunol. 2000, 18, 767–811; b) M. Cabeza-Cabrerizo, A. Cardoso, C. M. Minutti, M. Pereira Da Costa, C. Reis E Sousa, Annu. Rev. Immunol. 2021, 39, 131–166.

[4] a) G. Elia, Proteomics 2008, 8, 4012–4024; b) B. Wollscheid, D. Bausch-Fluck, C. Henderson, R. O’Brien, M. Bibel, R. Schiess, R. Aebersold, J. D. Watts, Nat. Biotechnol. 2009, 27, 378–386; c) Y. Li, Y. Wang, J. Mao, Y. Yao, K. Wang, Q. Qiao, Z. Fang, M. Ye, J. Proteomics 2019, 196, 33–41.

[5] a) Y. Liu, Y. Ge, R. Zeng, W. S. C. Ngai, X. Fan, P. R. Chen, CCS Chemistry 2023, 5, 802–813; b) W. Qin, K. F. Cho, P. E. Cavanagh, A. Y. Ting, Nature Methods 2021, 18, 133–143; c) L. L. Kirkemo, S. K. Elledge, J. Yang, J. Byrnes, J. Glasgow, R. Blelloch, J. A. Wells, eLife 2022, 11; d) J. Li, S. Han, H. Li, N. D. Udeshi, T. Svinkina, D. R. Mani, C. Xu, R. Guajardo, Q. Xie, T. Li, D. J. Luginbuhl, B. Wu, C. N. McLaughlin, A. Xie, P. Kaewsapsak, S. R. Quake, S. A. Carr, A. Y. Ting, L. Luo, Cell 2020, 180, 373-386.e315; e) K. H. Loh, P. S. Stawski, A. S. Draycott, N. D. Udeshi, E. K. Lehrman, D. K. Wilton, T. Svinkina, T. J. Deerinck, M. H. Ellisman, B. Stevens, S. A. Carr, A. Y. Ting, Cell 2016, 166, 1295-1307.e1221; f) Y. Li, Y. Wang, Y. Yao, J. Lyu, Q. Qiao, J. Mao, Z. Xu, M. Ye, Anal. Chem. 2021, 93, 4542–4551; g) S. A. Shuster, J. Li, U. Chon, M. C. Sinantha-Hu, D. J. Luginbuhl, N. D. Udeshi, D. K. Carey, Y. H. Takeo, Q. Xie, C. Xu, D. R. Mani, S. Han, A. Y. Ting, S. A. Carr, L. Luo, Neuron 2022, S0896627322008649.

[6] N. D. Udeshi, K. Pedram, T. Svinkina, S. Fereshetian, S. A. Myers, O. Aygun, K. Krug, K. Clauser, D. Ryan, T. Ast, V. K. Mootha, A. Y. Ting, S. A. Carr, Nat. Methods 2017, 14, 1167–1170.

[7] a) J. M. R. Narayanam, C. R. J. Stephenson, Chem. Soc. Rev. 2011, 40, 102–113; b) J. Zhao, W. Wu, J. Sun, S. Guo, Chem. Soc. Rev. 2013, 42, 5323; c) J. Twilton, C. C. Le, P. Zhang, M. H. Shaw, R. W. Evans, D. W. C. MacMillan, Nat. Rev. Chem. 2017, 1, 1–19; d) A. Y. Chan, I. B. Perry, N. B. Bissonnette, B. F. Buksh, G. A. Edwards, L. I. Frye, O. L. Garry, M. N. Lavagnino, B. X. Li, Y. Liang, E. Mao, A. Millet, J. V. Oakley, N. L. Reed, H. A. Sakai, C. P. Seath, D. W. C. MacMillan, Chem. Rev. 2022, 122, 1485–1542.

[8] S. Qiu, W. Li, T. Deng, A. Bi, Y. Yang, X. Jiang, J. P. Li, Angew. Chem. Int. Ed. 2023, 62, e202303014.

[9] S. H. Møller, L. Wang, P.-C. Ho, Cell. Mol. Immunol. 2021, 19, 370–383.

[10] M. Merad, P. Sathe, J. Helft, J. Miller, A. Mortha, Annu. Rev. Immunol. 2013, 31, 563–604.

[11] a) H. Donnelly, E. Mandrou, R. Insall, Sci. Immunol. 2023, 8, eadj3102; b) R. Chen, L. Chen, Trends Cell Biol. 2022, 32, 186–201.

[12] L. Becker, N.-C. Liu, M. M. Averill, W. Yuan, N. Pamir, Y. Peng, A. D. Irwin, X. Fu, K. E. Bornfeldt, J. W. Heinecke, PLoS ONE 2012, 7.

[13] a) M. B. Lutz, N. Kukutsch, A. L. J. Ogilvie, S. Rößner, F. Koch, N. Romani, G. Schuler, J. Immunol. Methods 1999, 223, 77–92; b) M. B. Lutz, S. Ali, C. Audiger, S. E. Autenrieth, L. Berod, V. Bigley, L. Cyran, M. Dalod, J. Dörrie, D. Dudziak, G. Flórez-Grau, L. Giusiano, G. J. Godoy, M. Heuer, A. B. Krug, C. H. K. Lehmann, C. T. Mayer, S. H. Naik, S. Scheu, G. Schreibelt, E. Segura, K. Seré, T. Sparwasser, J. Tel, H. Xu, M. Zenke, Eur. J. Immunol. 2022, eji.202249816.

[14] a) K. Ni, H. O’Neill, Immunol. Cell Biol. 1997, 75, 223–230; b) C. Hivroz, K. Chemin, M. Tourret, A. Bohineust, Crit. Rev. Immunol 2012, 32, 139–155; c) A. Chudnovskiy, G. Pasqual, G. D. Victora, Curr. Opin. Immunol. 2019, 58, 24–30.

[15] a) M. E. Futschik, B. Carlisle, J. Bioinform. Comput. Biol. 2005, 03, 965–988; b) L. Kumar, M. E. Futschik, Bioinformation 2007, 2, 5–7.

[16] a) C. D. Platt, J. K. Ma, C. Chalouni, M. Ebersold, H. Bou-Reslan, R. A. D. Carano, I. Mellman, L. Delamarre, Proc. Natl. Acad. Sci. 2010, 107, 4287–4292; b) O. Takeuchi, S. Akira, Cell 2010, 140, 805–820; c) W.-M. Chu, Cancer Lett. 2013, 328, 222–225.

[17] J. Liu, X. Zhang, Y. Cheng, X. Cao, Cell. Mol. Immunol. 2021, 18, 2461–2471.

[18] K. Liu, G. D. Victora, T. A. Schwickert, P. Guermonprez, M. M. Meredith, K. Yao, F.-F. Chu, G. J. Randolph, A. Y. Rudensky, M. Nussenzweig, Science 2009, 324, 392–397.

[19] a) E. P. Brandum, A. S. Jørgensen, M. M. Rosenkilde, G. M. Hjortø, Int. J. Mol. Sci. 2021, 22, 8340; b) F. Jimenez, M. P. Quinones, H. G. Martinez, C. A. Estrada, K. Clark, E. Garavito, J. Ibarra, P. C. Melby, S. S. Ahuja, J. Immunol. 2010, 184, 5571–5581; c) B. C. Urban, N. Willcox, D. J. Roberts, Proc. Natl. Acad. Sci. 2001, 98, 8750–8755; d) Y. Chen, J. Zhang, W. Cui, R. L. Silverstein, J. Exp Exp. 2022, 219, e20211314.

[20] J. I. Rodriguez-Barbosa, P. Schneider, A. Weigert, K.-M. Lee, T.-J. Kim, J.-A. Perez-Simon, M.-L. Del Rio, Cell. Mol. Immunol. 2019, 16, 679–682.

[21] T. B. H. Geijtenbeek, S. I. Gringhuis, Nat. Rev. Immunol. 2009, 9, 465–479.

[22] J. Helft, J. Böttcher, P. Chakravarty, S. Zelenay, J. Huotari, Barbara U. Schraml, D. Goubau, C. Reis e Sousa, Immunity 2015, 42, 1197–1211.

[23] Y. Shapir Itai, O. Barboy, R. Salomon, A. Bercovich, K. Xie, E. Winter, T. Shami, Z. Porat, N. Erez, A. Tanay, I. Amit, R. Dahan, Cell 2024, 187, 375-389.e318.

[24] H.-W. Rhee, P. Zou, N. D. Udeshi, J. D. Martell, V. K. Mootha, S. A. Carr, A. Y. Ting, Science 2013, 339, 1328–1331.

[25] T. Wu, E. Hu, S. Xu, M. Chen, P. Guo, Z. Dai, T. Feng, L. Zhou, W. Tang, L. Zhan, X. Fu, S. Liu, X. Bo, G. Yu, The Innovation 2021, 2, 100141.

[26] a) T. Chen, J. Ma, Y. Liu, Z. Chen, N. Xiao, Y. Lu, Y. Fu, C. Yang, M. Li, S. Wu, X. Wang, D. Li, F. He, H. Hermjakob, Y. Zhu, Nucleic Acids Res. 2022, 50, D1522–D1527; b) J. Ma, T. Chen, S. Wu, C. Yang, M. Bai, K. Shu, K. Li, G. Zhang, Z. Jin, F. He, H. Hermjakob, Y. Zhu, Nucleic Acids Res. 2019, 47, D1211–D1217.

