## Supplementary Figures. for "Cocktail Chemical Labeling for In-Depth Surfaceome Profiling of Bone-Marrow Derived Dendritic Cells"

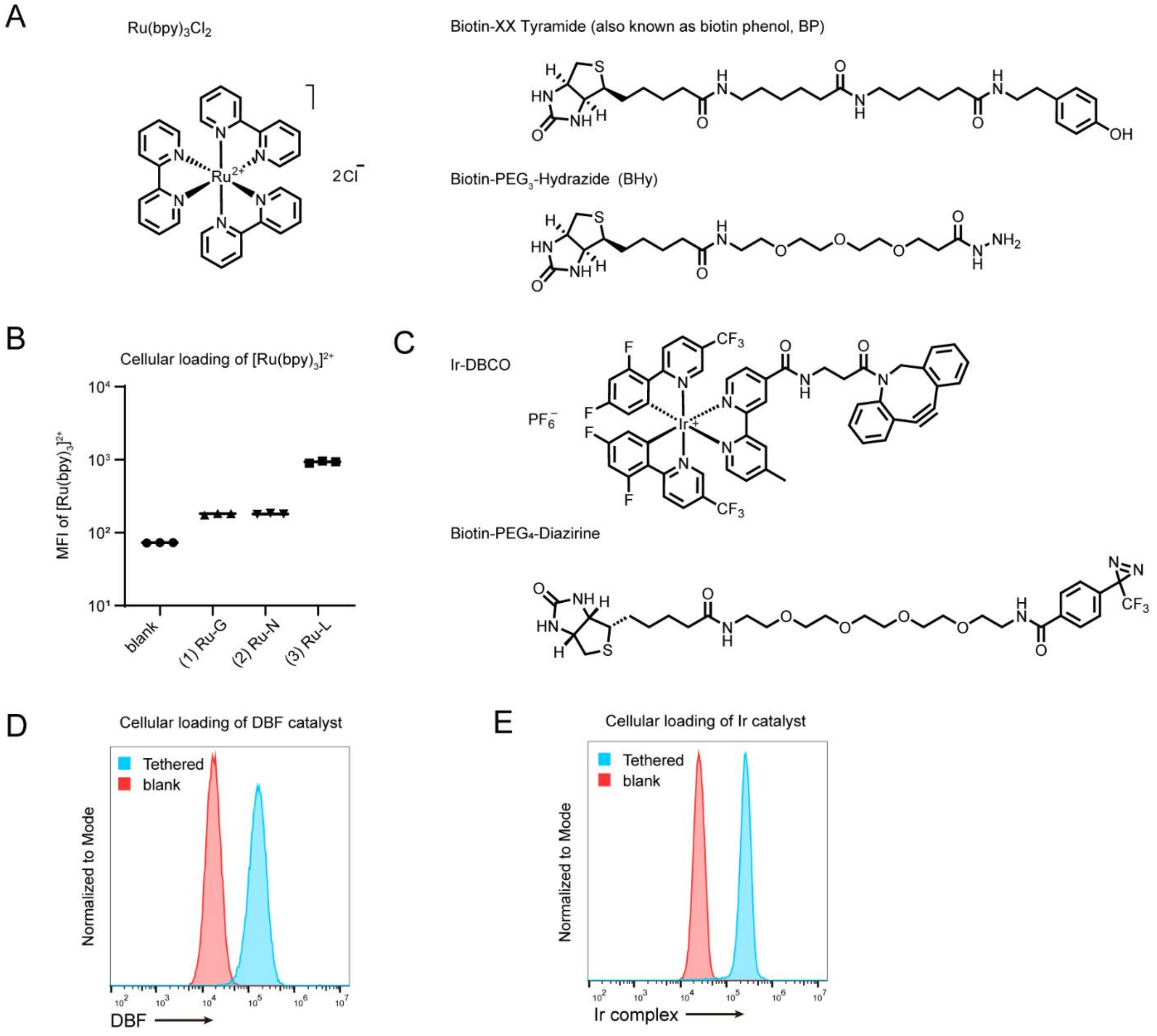


**Figure S1.** Evaluation of cell membrane-tethering effects on different photoredox catalysts. (A) Structure of the reagents in the Ru complex labeling toolkit. (B) Loading of cell-tethering Ru derivatives, as indicated by the emission at the PerCP-Cy5.5 channel. Data are represented as mean±S.D. (n=3). The loading amounts, in ascending order, are as follows: Ru-G, Ru-N, Ru-L. (C) Structure of the reagents used in the Ir complex labeling toolkit. Representative flow cytometric results of (D) DBF catalyst and (E) Ir catalyst at cell surface measured at the FITC channel and AmCyan channel, respectively.

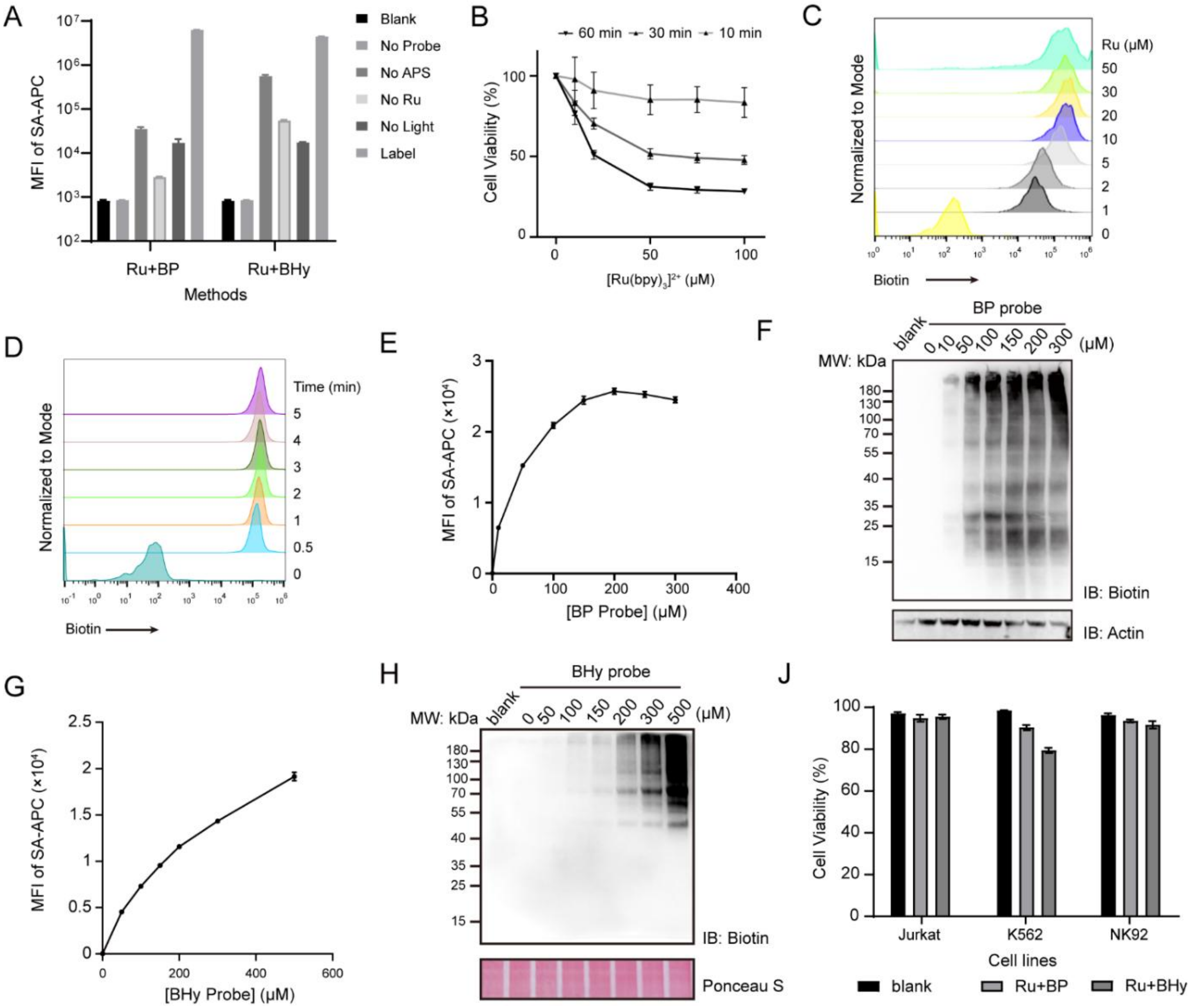


**Figure S2**. Optimization of cell surface labeling mediated by free-Ru. (A) Control experiments for Ru-mediated photo-reaction. Optimization of free Ru concentrations based on (B) the cytotoxicity of Ru-mediated PPL in the absence of biotin probes and (C) labeling efficiency using the BP probe. Time in (B) indicates the post-treatment culture time after irradiation in the presence of only Ru and APS. Cell viabilities remained above 80 % after a 10-min post-treatment culture. (D) Optimization of irradiation time using a 450 nm light source. Optimization of BP/BHy probe concentration as determined by (E, G) flow cytometry analysis and (F, H) western blot. (J) Verification of the biocompatibility of the Ru-mediated PPL reaction using three immune cell lines: Jurkat, K562, and NK92. All the experiments in this figure except (J) were performed on HeLa cells. Data are represented as mean±S.D. (n=3)

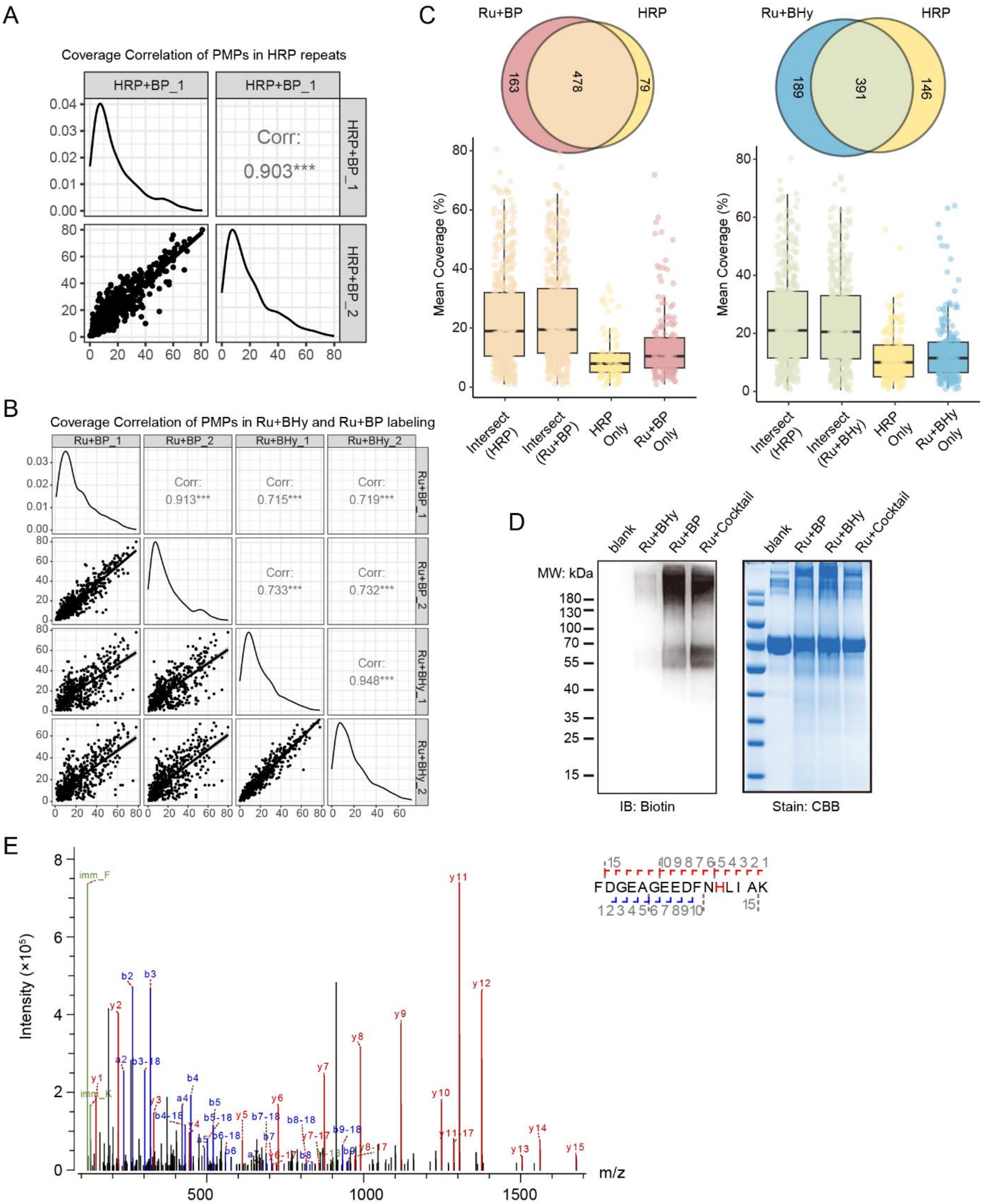


**Figure S3**. Differential labeling substrate scope of PECSL, Ru+BP, and Ru+BHy techniques revealed by LC-MS/MS analysis. Verification of the reproducibility of proteomic experiments following (A) the PECSL approach or (B) Ru+BP/BHy approach by correlation of sequence coverage. (C) Venn diagrams representing proteins commonly detected in two biological replicates involving either PECSL or Ru+BP/BHy, along with the boxplot showing average sequence coverage of the corresponding protein sets. (D) Immunoblotting assays and Coomassie brilliant blue (CBB) staining confirmed successful biotinylation on the BSA model protein via Ru-mediated PPL. (E) Representative MS/MS spectra of peptides with dioxidized histidine residue of Ru+BP labeling approach treated *Cp*OGA$^{D298N}$ sample.

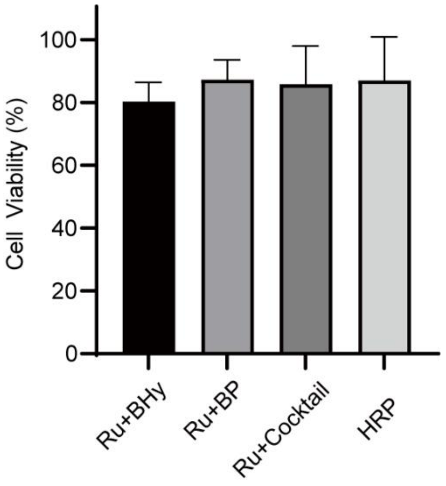


**Figure S4**. Normalized cell viability after three Ru-mediated labeling and the HRP-mediated labeling approaches.

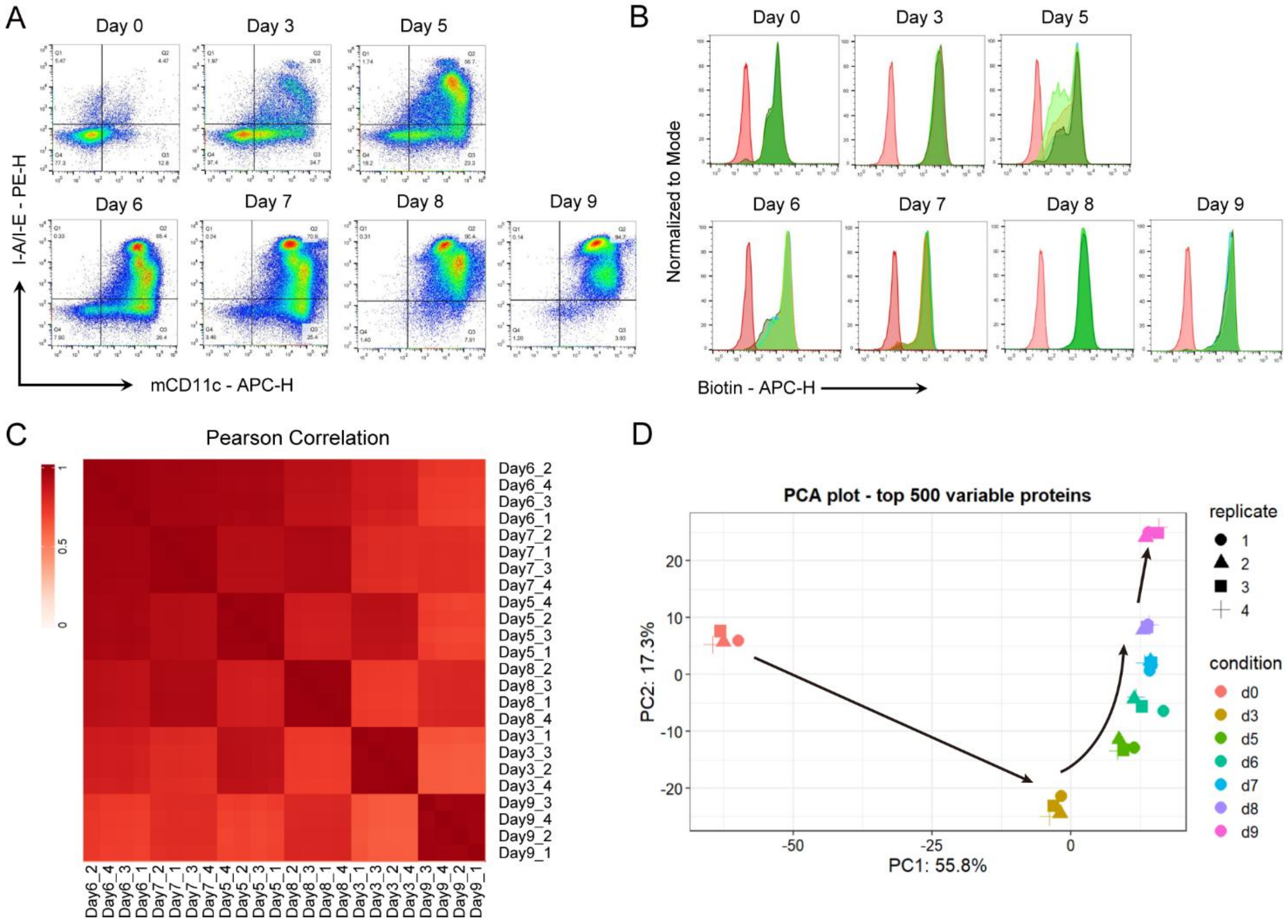


**Figure S5**. Quality control of the BMDC surfaceome profiling workflow. (A) Representative flow cytometry assays of the continuous BMDC maturation process. More than 90% of the cells in the plate were BMDCs after day 8, and the maturity of the BMDCs increased with LPS stimulation. (B) Cell surface biotinylation intensity of BMDCs harvested at different days during the differentiation process measured by flow cytometry. (C) Heat map showing all pairwise Pearson correlations among the 24 samples (4 replicates for each selected day) for MaxLFQ determined protein abundance values. (D) Principal component analysis of protein abundance data reflecting the BMDCs corresponding to different time points.

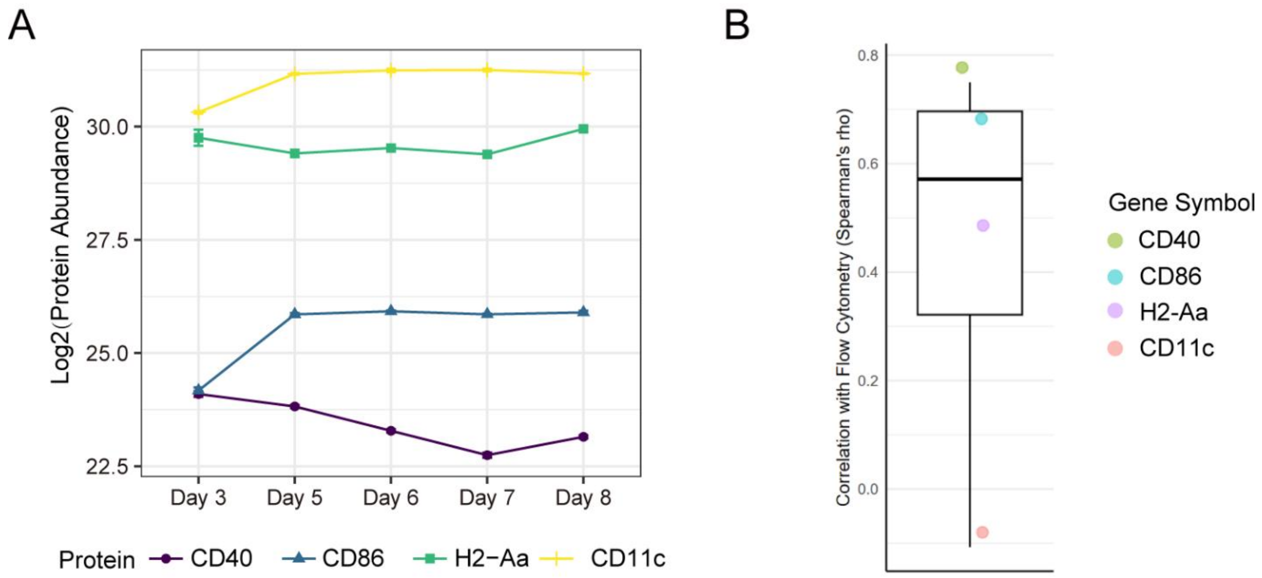


**Figure S6**. Mass spectrometric quantification and correlation analysis of BMDC maturity markers. (A) Mass spectrometric quantification of the four classic markers for characterization of BMDC maturity: CD40, CD86, MHC II (marked as H2-Aa), CD11c. (B) Correlation analysis of selected genes by Spearman's rho between mass spectrometric quantification and flow cytometry.
